# PepCCD: A Contrastive Conditioned Diffusion Framework for Target-Specific Peptide Generation

**DOI:** 10.1101/2025.09.01.673427

**Authors:** Jun Zhang, Yangyang Zhou, Tiantian Zhu, Zexuan Zhu

## Abstract

Peptide-based drug design targeting “undruggable” proteins remains one of the most critical challenges in modern drug discovery. Conventional peptide-discovery pipelines rely on low-throughput experimental screening, which is both time-consuming and prohibitively expensive. Moreover, existing computational approaches for designing peptides against target proteins typically depend on the availability of high-quality structural information. Although recent structure-prediction tools such as AlphaFold3 have achieved breakthroughs in protein modeling, their accuracy for functional interfaces remains limited. The acquisition of high-resolution structures is often expensive, time-intensive, and particularly challenging for targets with dynamic conformations, further restricting the efficient development of peptide therapeutics. Additionally, current sequence-based generative methods follow a paradigm that relies on known templates, which limits the exploration of sequence space and results in generated peptides lacking diversity and novelty. To address these limitations, we propose a contrastive conditioned diffusion framework for target-specific peptide generation, referred to as PepCCD. It employs a contrastive learning strategy between proteins and peptides to extract sequence-based conditioning representations of target proteins, which serve as precise conditions to guide a pre-trained diffusion model to generate peptide sequences with the desired target specificity. Extensive experiments on multiple benchmark target proteins demonstrate that the peptides designed by PepCCD exhibit strong binding affinity and outperform state-of-the-art methods in terms of diversity and generation efficiency.

## Introduction

In modern drug development, more than 80% of pathogenic proteins are considered “undruggable” by traditional small-molecule inhibitors due to the lack of stable binding pockets (Dang et al. 2017; Behan et al. 2019). As a therapeutic modality positioned between small molecules and antibodies, peptide-based drugs have emerged as promising candidates to target these undruggable proteins, due to their favorable target specificity, biocompatibility, and low toxicity profiles (Bodanszky 1988; Craik et al. 2013; Fosgerau and Hoffmann 2015a; Gomes et al. 2018; Muttenthaler et al. 2021a). Peptides can achieve precise modulation of disease-related targets through mechanisms such as disrupting protein–protein interactions (PPI), inhibiting enzymatic activity, or inducing targeted protein degradation (Fosgerau and Hoffmann 2015b). To date, over 100 peptide drugs have been approved for the treatment of various diseases, including cancer, diabetes, and HIV (Kaspar and Reichert 2013; Henninot, Collins, and Nuss 2018; Lee et al. 2019; Wang et al. 2022; Chen et al. 2024b). Traditionally, the discovery of therapeutic peptides has relied on high-throughput techniques such as yeast display (Muttenthaler et al. 2021b) or computational tools tailored to specific peptide properties (Lee et al. 2017; Lee, Wong, and Ferguson 2018) to explore the vast sequence space. However, the combinatorial space of potential peptides is huge, and only a small subset of sequences satisfies therapeutic requirements, making brute-force screening approaches both time-consuming and expensive.

To overcome the limitations of traditional brute-force screening, increasing attention has turned to peptide drug design guided by specific information about the target protein. However, designing peptides for specific targets remains a highly challenging task. First, conventional computational design methods (Wang et al. 2024; Watson et al. 2023; Anishchenko et al. 2021) heavily rely on high-resolution three-dimensional protein structures. Although recent deep learning-based structure prediction tools, such as AlphaFold3 (Abramson et al. 2024), have achieved remarkable breakthroughs, they still suffer from limited accuracy in predicting critical binding interfaces and modeling proteins with multiple conformational states (Pereira et al. 2025; Chakravarty, Lee, and Porter 2025). High-quality structural data are still challenging to obtain in practice, especially for intrinsically disordered proteins that lack stable conformations (Oldfield and Dunker 2014; Maiti et al. 2024; Trivedi and Nagarajaram 2022). Second, the peptide sequence space is vast, making the design and search process highly challenging (Jenson et al. 2018). Recently, several innovative sequence-only design approaches have been proposed (Bhat et al. 2025; Chen et al. 2024a); however, these computational methods typically rely heavily on initial template sequences, and the scarcity of high-quality peptide–protein binding data further limits the exploration of the sequence space. As a result, the generated peptides often lack diversity and novelty.

Motivated by the above challenges, we propose a **C**ontrastive **C**onditioned **D**iffusion framework for target-specific peptide generation, named **PepCCD**. Unlike existing sequence-guided approaches, PepCCD adopts a generative modeling paradigm, directly using the target protein sequence as a conditioning input to guide the generation of peptide sequences. Overall, the main contributions of this work are as follows:

- We are the first to introduce diffusion models for the design of peptides guided by target protein sequences. PepCCD directly trains a conditional diffusion model, in which target protein sequences are encoded as condition vectors using a contrastive learning framework. The model generates peptide sequences in an end-to-end manner without the need to construct candidate peptide libraries or perform post hoc screening.
- We construct a large-scale synthetic peptide fragment dataset to pre-train the conditional diffusion model. This dataset enhances the model’s prior knowledge of peptide sequences and compensates for the scarcity of experimentally determined protein-peptide complex data, thereby improving the diversity of the generated peptides.
- Extensive experiments on benchmark target proteins demonstrate that PepCCD outperforms current state-of-the-art sequence-guided baselines in generating target-specific peptides, particularly excelling in terms of targeting accuracy, diversity, and generation efficiency.

## Related Works

### Protein Language Models

Recent advances in deep learning–based protein sequence modeling have opened new avenues for peptide drug discovery. Protein language models (pLMs) pre-trained on large-scale protein sequence databases can now capture rich semantic, structural, and functional patterns, enabling structure-free peptide design (Rives et al. 2021; Shin et al. 2021). These breakthroughs have driven significant progress in two key areas: (1) substantial improvements in the effectiveness of protein sequence embeddings (Hayes et al. 2025; Brandes et al. 2022; Lin et al. 2023); and (2) strong adaptability to downstream tasks such as protein design after fine-tuning on task-specific data (Madani et al. 2023; Ferruz, Schmidt, and Höcker 2022; Lv et al. 2025). Inspired by these advances, this study leverages protein language models fine-tuned specifically for peptide generation, enabling *de novo* design of target-specific peptides using only the target protein sequence.

### Diffusion Models

Diffusion models, which generate new samples by gradually adding noise to input data and learning to reverse the process from a prior distribution, have emerged as a robust generative framework across various domains, including image and text synthesis (Song and Ermon 2019; Trippe et al. 2022). Recently, their potential has been explored in protein design. For example, Liu et al. (Liu et al. 2025) introduced a text-conditional diffusion model to generate proteins based on natural language prompts. Hoogeboom et al.

(Hoogeboom et al. 2022) developed ProtDiff, which leverages E(3)-equivariant graph neural networks to model the diverse distributions of protein backbone coordinates. Despite these advances, the integration of diffusion models into peptide–protein sequence design remains largely unexplored. In particular, no prior work has addressed target protein sequence–guided peptide generation using diffusion models. This study presents the first diffusion-based framework conditioned on protein sequences for *de novo* design of target-specific peptides, filling a critical gap in the field.

### Contrastive Learning

As a widely used approach in self-supervised learning, contrastive learning enables models to capture hidden patterns in data without requiring explicit labels. Its core principle is to optimize the interaction of samples in the embedding space, pulling positive pairs closer while pushing negative pairs farther apart. Yuan et al. (Yuan et al. 2021) proposed a multimodal contrastive learning framework to align textual and visual data by enhancing the similarity of related text–image pairs and suppressing the similarity of unrelated ones, thereby achieving cross-modal alignment. Zhang et al. (Zhang et al. 2023) applied conformational augmentation to protein structures, thereby maximizing the similarity between embeddings of the same protein and the dissimilarity between different proteins, which enables more discriminative representations. Inspired by these contrastive learning strategies, we design a protein–peptide sequence contrastive learning framework that extracts informative features from protein sequences and uses them as conditional inputs to guide the generation of target-specific peptides.

## Methodology

### Datasets

To support both pre-training and fine-tuning of the model, we constructed two complementary datasets: a protein–peptide interaction dataset (S1) for fine-tuning, and a large-scale simulated peptide dataset (S2) for pre-training. The construction of each dataset is described as follows.

#### Fine-tuning Dataset S1

To train the model to recognize and generate peptides with target-binding capabilities, we followed the PepPrCLIP protocol and constructed a high-quality protein–peptide interaction dataset based on co-crystal structures from the RCSB Protein Data Bank. We defined valid interactions as those with a buried surface area (BSA) greater than or equal to 50 square angstroms. This dataset includes a wide range of binding modes, ranging from strong to weak affinities, effectively capturing diverse real-world interactions. To ensure data diversity and reduce redundancy, we clustered all protein sequences using CD-HIT (Fu et al. 2012) with a sequence identity threshold of 0.9. The final dataset consists of 15,110 complex pairs for training and 5,480 for testing.

#### Pre-training Dataset S2

Inspired by transfer learning strategies, we constructed a simulated peptide pre-training dataset based on UniProt to enhance the model’s generalization to peptide sequences and provide it with broad prior knowledge. We applied a sliding window technique, with a window size of 30 amino acids and a stride of 15 amino acids, to segment sequences longer than a specified threshold. Sequences containing non-canonical amino acids (B, J, O, U, X, Z) were removed. This process generated a total of 68,958,049 simulated peptide sequences that were shorter than 30 amino acids in length. The sliding window approach is widely used in peptide function prediction and screening tasks, and its effectiveness and validity in sequence modeling have been demonstrated in previous studies.

### Overview of PepCCD

The overall framework of the proposed method is illustrated in Figure 1(A), where a potential target-specific peptide can be designed based solely on the input target protein sequence. The protein encoder is a pre-trained ESM-2 protein language model, fine-tuned via protein–peptide semantic alignment, and is used to extract features from the protein sequence as a conditional vector to guide peptide generation. By sampling Gaussian noise to obtain hidden states, which are then concatenated with the target condition vector to serve as the input to the conditional diffusion model. The model then denoises this input to produce a latent representation, which is subsequently decoded into the final peptide sequence. In this study, we introduce a three-stage training strategy that progressively enhances the model’s ability to generate peptides with both diversity and target specificity.

**Figure 1:**
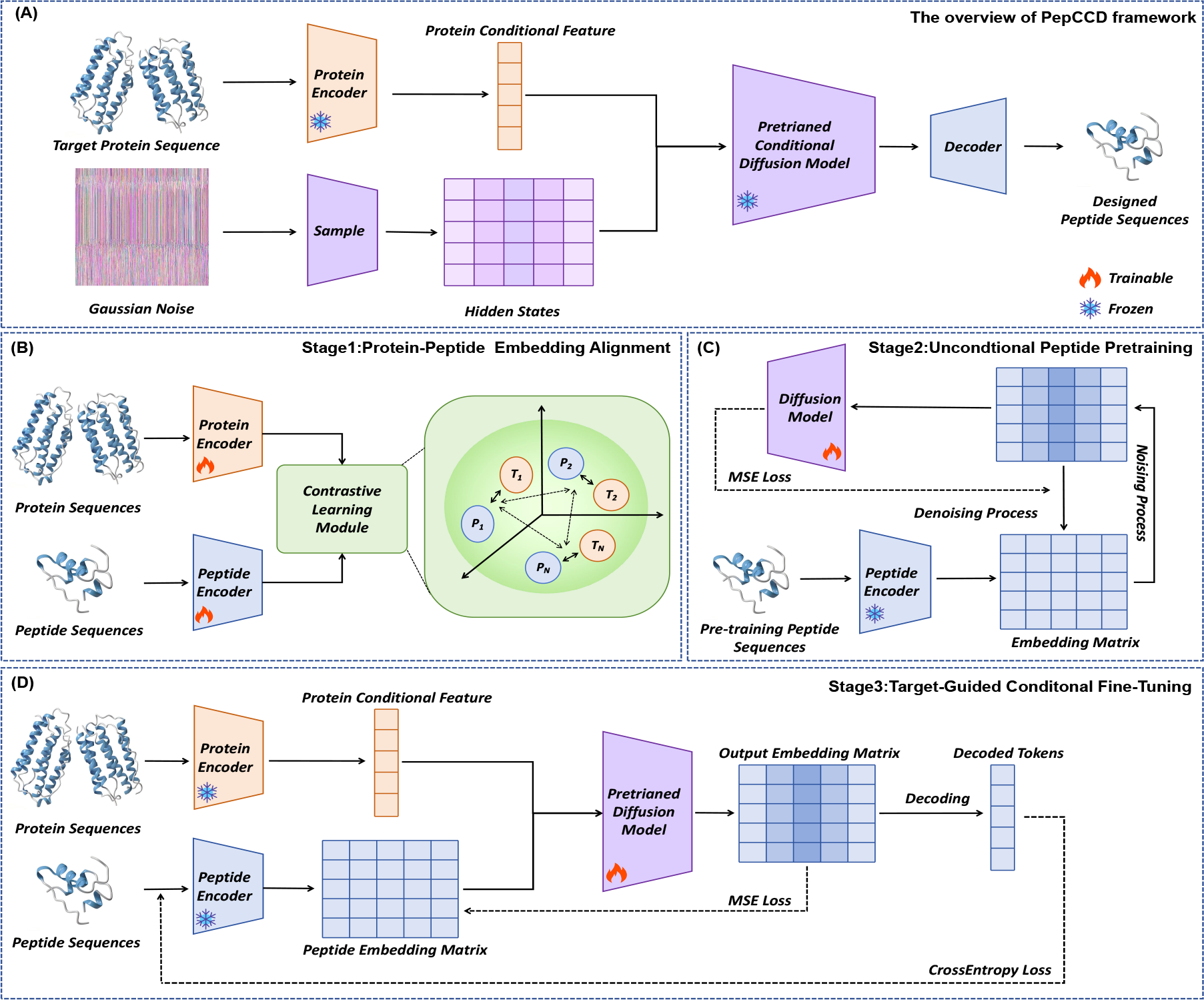
Overview of the PepCCD framework. (A) The overall architecture for generating target-specific peptides from protein sequences. (B) Protein–peptide semantic alignment via contrastive learning using dual ESM-2 encoders. (C) Unconditional pretraining of the diffusion model on unlabeled peptides. (D) Target-guided fine-tuning for peptide generation using conditional denoising.

### Stage 1: Protein-Peptide Alignment

Inferring latent peptides solely from a protein sequence is a challenging task that requires identifying the correct protein–peptide pairing within a many-to-many mapping. To establish a semantic bridge between proteins and peptides, we introduce a joint embedding framework built on contrastive learning. To capture deep sequential and structural semantics while avoiding discrepancies arising from different protein language models, we employ two identical ESM-2 encoders—one for protein sequences and one for peptide sequences. ESM-2 is a large-scale pre-trained protein language model that encodes rich structural, functional, and evolutionary information embedded in amino-acid sequences, and has proven effective across diverse downstream tasks. This property enables PepCCD to design structurally meaningful peptides even in the absence of target protein structures.

Figure 1(B) illustrates the semantic alignment process. During training, both encoders are fine-tuned on the constructed dataset S1. The training objective is to use contrastive learning to ensure that matched protein–peptide pairs exhibit high similarity in the feature space, while unmatched pairs exhibit low similarity. In this stage, we employ the InfoNCE loss to achieve this goal, formulated as follows:

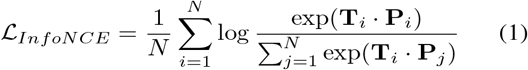

Here, **T**_*i*_ and **P**_*i*_ represent the normalized embedding vectors of the target protein and peptide, respectively. Essentially, the objective is to maximize the probability of the positive pair among all possible candidates. In this stage, we evaluated the performance of different types of encoders for the protein–peptide semantic alignment task. The detailed results are shown in the Appendix.

### Stage 2: Unconditional Peptide Pre-training

In Stage 1, we obtained a pair of encoders that can accurately align and capture protein–peptide semantic relationships. To further enhance the model’s stability and improve the diversity of peptide generation, we introduce a largescale unlabeled peptide dataset S2 in this stage for the unconditional pre-training of the diffusion model. The diffusion model comprises two primary processes: a noise addition process and a denoising process. In the noise addition phase, peptide embeddings undergo a forward diffusion transformation, gradually injected with Gaussian noise. In the denoising phase, a Transformer-based architecture is used to reconstruct the embeddings. This Transformer consists of multiple modules, each incorporating multilayer perceptrons (MLPs) and multi-head self-attention, enabling the model to capture complex noise patterns by modeling dependencies across sequence positions.

As shown in Figure 1(C), we first use the pre-trained peptide encoder to extract embedding representations from the unlabeled peptide sequences, allowing the model to adapt to the target-conditioned generation task. Gaussian noise is then added to the embeddings over 500 time steps, simulating the diffusion process as a Markov chain. The diffusion model learns to reconstruct the original embeddings from the noisy representations, thereby capturing the prior distribution of peptide features.

At this stage, we utilize the Mean Squared Error (MSE) as the loss function. The training objective is to constrain the diffusion model to minimize the discrepancy between the reconstructed and original embeddings, enabling it to effectively learn the complex distribution within the peptide embedding space and acquire rich semantic representations. The loss function is defined as follows:

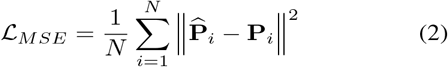

where **P**_*i*_ and 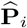 denote the original embedding matrix extracted by the pre-trained peptide encoder and the embedding matrix reconstructed from noise by the diffusion model, respectively.

### Stage 3: Target-Guided Conditioned Fine-Tuning

Following the first two stages, we use the pre-trained encoders to extract the target protein condition vectors and peptide embedding matrices from the fine-tuning dataset. The condition vectors serve as guidance for the pre-trained diffusion model to perform denoising and generate potential target-specific peptide sequences.

As illustrated in Figure 1(D), the primary objective of this stage is to adapt the diffusion model for target-specific generation further. Unlike the pre-training phase, the diffusion model now takes both Gaussian noise and the target condition vector as input to reconstruct the peptide embedding matrix. The reconstructed embeddings are then decoded into amino acid sequences, and a cross-entropy loss is applied to measure the discrepancy between the generated peptides and ground truth sequences.

The goal of this stage is to minimize the cross-entropy loss between generated and actual peptide sequences, thereby improving the model’s accuracy and biological relevance. The cross-entropy loss function is defined as follows:

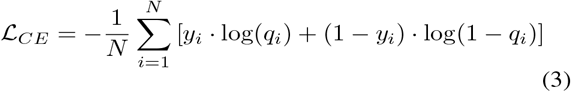

where *y*_*i*_ denotes the label of the *i*-th amino acid, which is equivalent to the token value of the input sequence, and *q*_*i*_ represents the output probability distribution of the amino acid at position *i*. Therefore, the fine-tuning stage is a multi-objective optimization process, and the complete loss function is shown below:

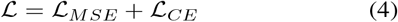

where ℒ_*MSE*_ denotes the reconstruction loss of the peptide embedding matrix, and ℒ_*CE*_ represents the decoding loss of the peptide sequence. This jointly optimized fine-tuning strategy significantly enhances the model’s capability to reconstruct complex peptide sequences, further consolidating the decoding performance of the denoising diffusion generative model. During the inference phase, Gaussian noise matrices and targeting conditions generated from extracted target protein sequences serve as guidance for random sampling, thereby generating target-binding peptides with rich diversity.

## Experiments

### Experiment Setup

#### Baselines

To evaluate the effectiveness of PepCCD, we compared it against two state-of-the-art approaches that represent distinct paradigms in peptide design: one that utilizes the structural information of the target protein (structure-guided), and another that conditions on the protein sequence (sequence-guided). These two methods—RFdiffusion and PepPrCLIP—are considered among the most competitive models for structure-based and sequence-based peptide generation, respectively.

It is worth noting that RFdiffusion was not initially designed for peptide sequence generation; rather, its primary focus lies in protein backbone modeling and structural design. However, due to its strong capabilities in structure generation, it has demonstrated promising potential in structure-guided peptide design tasks. Therefore, we adopt RFdiffusion as the representative structure-based method, and utilize ProteinMPNN(Dauparas et al. 2022) to generate amino acid sequences from its predicted backbones, following the same protocol as its official Colab implementation.

To ensure fairness and accuracy, we strictly followed the official implementation procedures of both RFdiffusion and PepPrCLIP, and deployed them in the local environment for consistent evaluation.

#### Evaluation Protocol

We conducted peptide drug design experiments on 209 target proteins, consistent with the evaluation set used by PepPrCLIP. For each target, every model was tasked with generating 10 novel peptide sequences. The quality of the generated peptides was assessed using the following metrics:

- **Interface TM score (ipTM)**: A key evaluation metric from the AlphaFold3 framework used to assess the binding affinity between the protein and peptide.
- **Rosetta-total-score(RT-score)** (Leaver-Fay et al. 2011): Measures the overall energy stability of the predicted complex, reflecting the overall stability of the complex interface by integrating various energy terms.
- **Sequence similarity**: Assesses the global sequence similarity between the generated peptide and its corresponding natural peptide, reflecting sequence-level diversity.
- **Structure similarity**: Quantifies the structural similarity between the generated and native peptides using the TM-score(Zhang et al. 2022), reflecting structural diversity.
- **Bioactivity**(Mooney et al. 2012): Predicts whether the generated peptide has potential biological activity, including functions such as antimicrobial effects or signal regulation.
- **Instability**(Gasteiger et al. 2005): Estimates the stability of the generated peptide under in vitro conditions, indicating its potential for experimental expression and application.

Detailed information on the tested target proteins, metrics, and implementation can be found in the Appendix. We only reported the average metrics over all generated peptides for each method in the experimental results.

## Experimental Results

### Performance Comparison

In the results of targeted peptide design performance, as shown in Table 3, PepCCD demonstrates significant advantages over a similar baseline method that is solely guided by sequence information, in terms of multiple key indicators evaluating the interaction between the peptide and target protein. This reflects its strong generative ability and modeling accuracy. Although PepCCD still slightly lags behind the state-of-the-art structure-guided model, RFDiffusion, which excels in predicting structured targets, this is primarily attributed to its direct utilization of protein three-dimensional structural information. Notably, all test targets in the benchmark are sourced from the PDB database and possess experimentally validated three-dimensional structures. It is plausible that these or structurally similar proteins were present in RFDiffusion’s training data, potentially offering it an advantage in these evaluations.

On the more generalizable metric—hit rate (calculated using ipTM(avg))—PepCCD even surpasses RFDiffusion, as shown in Figure 2. This demonstrates that, in settings where explicit structural information is unavailable and only sequence-based conditioning is used, PepCCD demonstrates exceptional stability and reliability in generating high-quality peptide candidates, ranking among the top-performing methods currently available.

**Figure 2:**
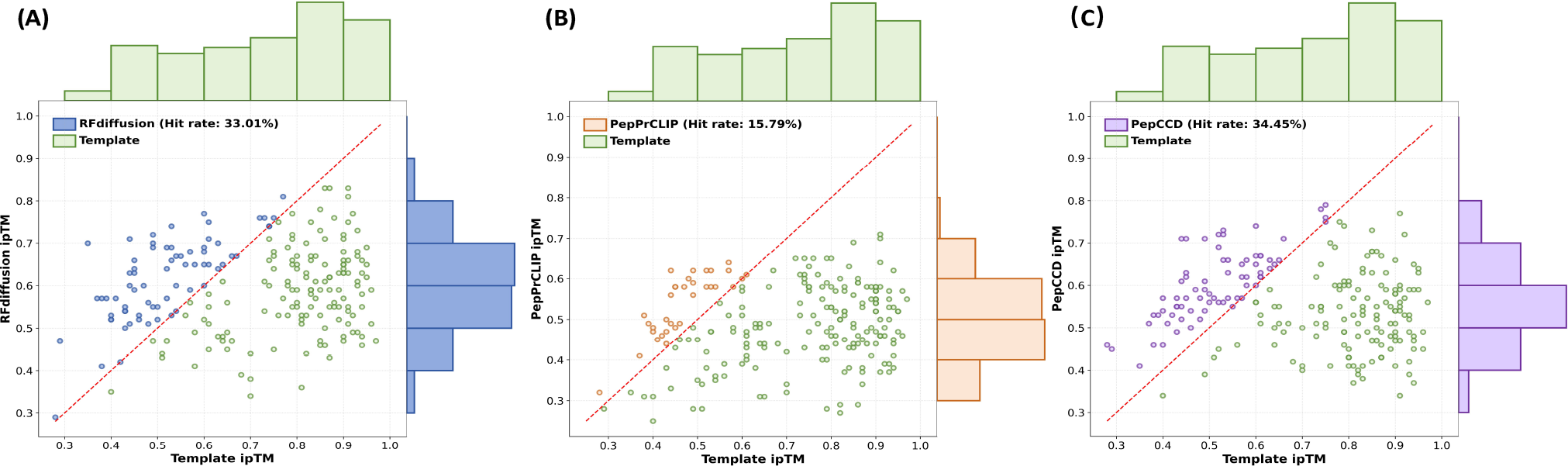
The hit rate is defined as the proportion of generated peptides whose ipTM(avg) scores are greater than or equal to those of their corresponding templates, serving as a metric to evaluate the model’s capability in producing high-quality peptide structures. Points above the diagonal line are considered hits. Comparisons of different models across various targets are presented in the Appendix.

In terms of overall performance and efficiency (as shown in Table 2), PepCCD achieves the lowest sequence and structural similarity, reduced instability, and the best bioactivity scores among all baseline methods, indicating strong generalization ability in generating diverse and stable peptide candidates. More importantly, PepCCD significantly outperforms existing complex structure-based modeling approaches in terms of inference efficiency, thereby greatly enhancing scalability in practical applications.

**Table 1:** Performance on target-oriented peptide design. In the column headers, **(best)** denotes the best single peptide designed for each target, whereas **(avg)** denotes the average performance across all targets.

| Methods | Type | ipTM(best)↑ | ipTM(avg)↑ | RT-score(best)↓ | RT-score(avg)↓ |
| --- | --- | --- | --- | --- | --- |
| RFdiffusion | Structure | **0.7838** | **0.5960** | **-394.422** | **-242.238** |
| PepPrCLIP | Sequence | 0.7141 | 0.4901 | -188.933 | -157.573 |
| PepCCD | Sequence | <u>0.7418</u> | <u>0.5557</u> | <u>-363.481</u> | <u>-188.933</u> |

**Table 2:**
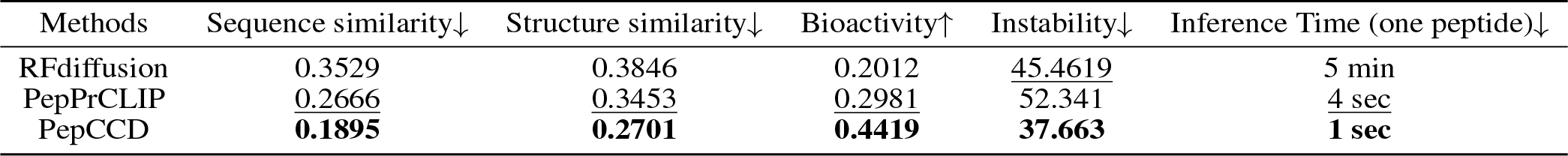
Overall Performance and Efficiency Comparison. Inference Time is the wall-clock time averaged over the full test set, measured on an NVIDIA GTX 4090 GPU.

| Methods | Sequence similarity↓ | Structure similarity↓ | Bioactivity↑ | Instability↓ | Inference Time (one peptide)↓ |
| --- | --- | --- | --- | --- | --- |
| RFdiffusion | 0.3529 | 0.3846 | 0.2012 | <u>45.4619</u> | 5 min |
| PepPrCLIP | <u>0.2666</u> | <u>0.3453</u> | <u>0.2981</u> | 52.341 | 4 sec |
| PepCCD | **0.1895** | **0.2701** | **0.4419** | **37.663** | **1 sec** |

We further evaluated the Global Amino-acid Composition Discrepancy **(GACD)**. As shown in Figure 3, peptides generated by PepCCD exhibit amino acid distributions closest to those of natural templates, indicating strong biological plausibility. In terms of overall performance and efficiency (as shown in Table 2), PepCCD achieves the lowest sequence and structural similarity, reduced instability, and the best bioactivity scores among all baseline methods, indicating strong generalization ability in generating diverse and stable peptide candidates. It also outperforms existing complex structure-based modeling approaches in terms of inference efficiency, thereby greatly enhancing scalability in practical applications.

**Figure 3:**
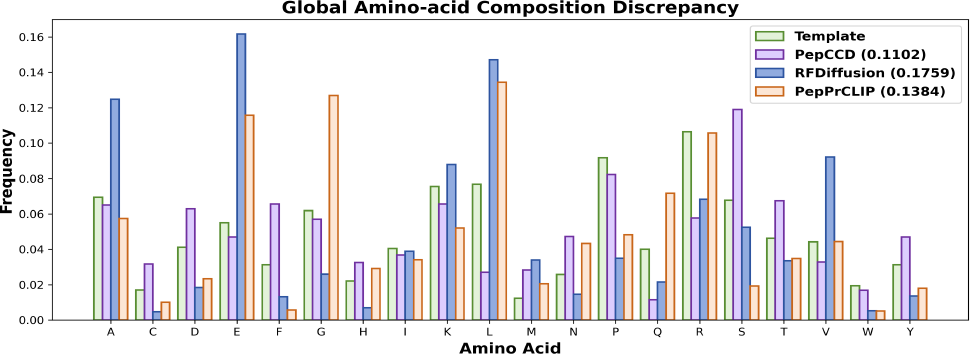
GACD is used to assess the difference in amino acid composition between peptide sequences and template peptides. The closer the frequency distribution is to the natural templates, the higher the biological relevance of the generated peptides will be. Detailed calculation procedures are provided in the Appendix.

### Ablation Study

To evaluate the necessity of each module, we compared PepCCD with three ablated variants: (1) PepCCD(w/o Align) that removes the align stage, (2) PepCCD(w/o Pre-training) that removes the pre-training stage, and (3)PepCCD(w/o Align & Pre-training) that removes both align and pre-training stages. The evaluation metrics used in the ablation study differ from those in the main comparison experiments. Since PepCCD is entirely sequence-based in both representation and generation, we avoided the high computational cost associated with AlphaFold3 complex structure prediction. Instead, we employed a more efficient sequence-level metric, **Superior ratio**, as a substitute for ipTM and RT-score (both of which rely on predicted structures and are computationally expensive). In addition, we introduced two new metrics — Intra Similarity **(Intra-Sim)** and Inter Similarity **(Inter-Sim)** — to evaluate the model’s performance in terms of intra-target diversity and inter-target specificity, respectively. The detailed calculation methods of these three new metrics are provided in the Appendix.

As shown in Table 3, removing the align module results in performance degradation across all metrics, with the Superior ratio exhibiting the most significant decline. Eliminating the pre-training module also results in an overall decline, particularly with a noticeable decrease in sequence diversity. When both modules are removed, the model’s performance deteriorates even further. Overall, the align and pretraining modules play essential roles in enhancing the specificity and diversity of the generated peptides, respectively. PepCCD achieves optimal performance when all modules are included, and the absence of any single component compromises both the biological plausibility and diversity of the generated peptide candidates.

**Table 3:**
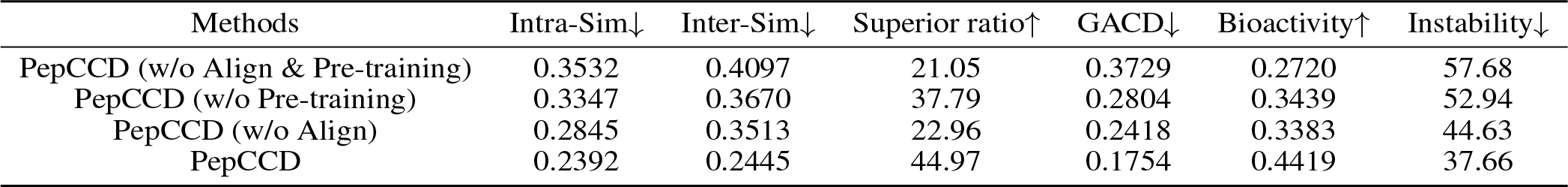
Ablation study results of PepCCD and its three variants across six evaluation metrics. Note that Intra Similarity measures the sequence similarity among peptides generated for the same target, reflecting diversity; Inter Similarity measures the similarity between peptides generated for different targets, reflecting the model’s ability to distinguish between targets; Superior ratio evaluates the target-specificity of the generated peptides.

### Molecular Dynamics Simulation

To further evaluate the binding affinity and structural stability of the designed peptides, we performed all-atom molecular dynamics (MD) simulations using the GROMACS software package (Abraham et al. 2015), and calculated binding free energies using MM/GBSA and MM/PBSA methods(Genheden and Ryde 2015). The simulations focused on interactions with the receptor-binding domain (RBD) of the SARS-CoV-2 spike protein (reference structure: PDB ID: 6M0J(Lan et al. 2020)). For each of the three methods, 10 representative peptides were selected. Their structures were predicted using AlphaFold3 and docked into the RBD binding pocket(Yan et al. 2020) via Rosetta. The most stable complex conformations were then subjected to standard preprocessing (Jo et al. 2008) and 100 ns production simulations to reflect physiological conditions. Full details of the simulation setup—including force field selection, solvation models, equilibration protocols, and free energy calculation parameters are provided in Appendix.

As shown in Figure 4, sequence-guided methods perform excellently in MD simulations, with peptides designed by PepCCD exhibiting the lowest binding free energies under both MM/GBSA and MM/PBSA calculations. Additionally, these peptides show significantly lower variance in binding energies, indicating better stability and consistency throughout the simulations. Notably, although RFDiffusion performs well on static target-binding metrics, it demonstrates the poorest performance in molecular dynamics simulations. This discrepancy may be due to the selected complex structure (PDB: 6M0J) lacking a native peptide ligand for the RBD, which prevents RFDiffusion from fully leveraging the advantages of its pre-trained structural information.

**Figure 4:**
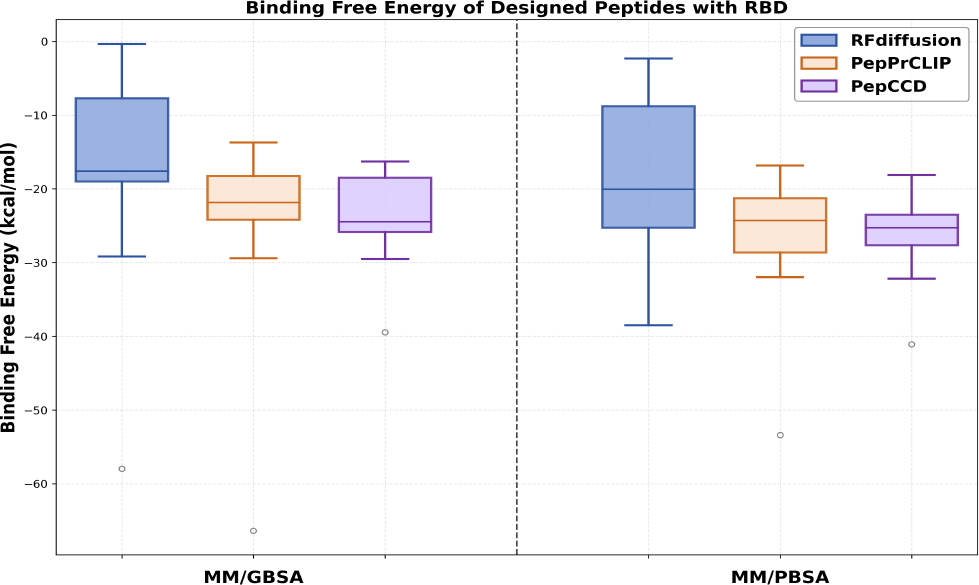
Based on the simulation trajectories, the binding free energies of each peptide–protein complex were calculated using the MM/GBSA and MM/PBSA methods, in order to assess the binding affinity between the peptides and the target protein.

## Conclusion

In this work, we introduced PepCCD, a novel contrastive conditioned diffusion framework designed for target-specific peptide generation based solely on protein sequence information. Unlike traditional approaches that rely on structural templates or limited sequence-guided heuristics, PepCCD integrates contrastive learning and diffusion-based generative modeling to achieve effective semantic alignment between proteins and peptides, thereby generating highly diverse, biologically relevant peptide sequences. Through a three-stage training paradigm—including protein–peptide alignment, large-scale unconditional peptide pre-training, and target-guided fine-tuning—PepCCD demonstrates superior performance in both generation quality and efficiency.

Extensive experiments on multiple benchmark datasets show that PepCCD outperforms existing sequence-based and even structure-based peptide generation methods in terms of binding affinity, diversity, bioactivity, and inference time. Furthermore, ablation studies validate the critical role of each component in enhancing specificity and diversity. At the same time, molecular dynamics simulations confirm the binding stability of peptides generated by PepCCD in realistic biological contexts. Overall, PepCCD offers a scalable and efficient structure-free solution for *de novo* peptide generation, especially valuable for targets lacking high-resolution structural data.

Thank you for reading these instructions carefully. We look forward to receiving your electronic files!

