## Appendix for "PepCCD: A Contrastive Conditioned Diffusion Framework for Target-Specific Peptide Generation"

### A:Encoder Performance for Alignment

To assess the impact of encoder architecture on the quality of semantic alignment between proteins and peptides, we conducted a comprehensive evaluation using several representative models. All models were trained using the same contrastive learning setup and **S1 dataset** to ensure a fair comparison. Specifically, we evaluated the following model variants:

- **ESM-2 (PepCCD proposed method)**(Lin et al. 2023): A protein language model developed by Meta AI based on the Transformer architecture, pre-trained on large-scale protein sequences. It effectively captures structural and functional information and is widely used in various downstream tasks.
- **ProtBERT** (Brandes et al. 2022): Measures the overall energy stability of the predicted complex, reflecting the overall stability of the complex interface by integrating various energy terms.
- **Transformer**: A Transformer model without pretraining, used as a baseline to evaluate the importance of pre-trained protein language models in the protein–peptide semantic alignment task.
- **nn.Embedding Encoder**: An encoder that directly embeds amino acid sequences using PyTorch's nn.Embedding layer, with fully random weights and no pretraining or contextual modeling, serving as a minimal performance baseline.

The results in Table2 highlight the critical role of encoder architecture in the protein–peptide semantic alignment task. Pretrained protein language models such as ESM-2 and ProtBERT significantly outperform non-pretrained models by better capturing deep semantic associations within sequences and improving both alignment and generation performance. ESM-2, in particular, demonstrates the strongest overall results, showcasing its advantage in modeling structural and functional semantics. In contrast, the randomly initialized Transformer shows a noticeable drop in performance, while the simple nn.Embedding encoder struggles to learn meaningful representations, reinforcing the indispensable role of pretraining and contextual modeling in protein–peptide alignment.

### B: Metrics for Model Comparison

For a fair and consistent comparison, each of the three models (RFdiffusion(Watson et al. 2023), PepPrCLIP(Bhat et al. 2025), and PepCCD) generated 10 peptide sequences for each target protein in the test set. For each target, we compute the average value of each evaluation metric over its generated peptides, representing the model's performance on that specific target. Finally, we aggregate the results across all targets and report the overall average for each metric as the model's final performance.

We selected six key metrics to comprehensively evaluate the quality of the generated peptides, covering aspects of structural biology, computational chemistry, and bioinformatics. Each metric is introduced with its evaluation purpose, followed by its specific computation method in this study. These metrics are listed below:

- **Interface TM-score (ipTM)**: This metric assesses the confidence of the predicted binding interface between the generated peptide and the target protein, reflecting their structural compatibility.In this study, we compute ipTM using AlphaFold3(Abramson et al. 2024), which predicts the 3D structure of the protein–peptide complex and outputs ipTM scores as confidence estimates for the predicted interfaces. Higher values (range: 0–1) indicate more confident binding.
- **Rosetta Total Score (RT-score)**: Used to evaluate the overall energetic stability of the protein–peptide complex, indirectly reflecting the biophysical plausibility of the binding conformation. In this study, we calculate this score using the **InterfaceAnalyzer**(Leaver-Fay et al. 2011) module of Rosetta, which considers factors such as van der Waals forces, solvation effects, and hydrogen bonding. Lower scores indicate more stable complex structures.
- **Sequence Similarity**: This metric measures the amino acid sequence similarity between the generated peptide and its corresponding native peptide, reflecting the trade-off between conservation and diversity. In this study, we perform global sequence alignment between each generated peptide and its matched native peptide using the **Bio.pairwise2** module.
- **Structural Similarity**: This metric quantifies the structural diversity of the generated peptides using the TM-score. In this study, we evaluate the predicted structures of the generated peptides against their native counterparts using **US-align**(Zhang et al. 2022). A lower TM-score generally indicates lower structural similarity, implying higher diversity.
- **Bioactivity**:This metric evaluates the potential biological functionality of a peptide, such as antimicrobial effects or immune regulation. In this study, we compute the bioactivity score using the **PeptideRanker**(Mooney et al. 2012), which predicts the likelihood of bioactivity based on the characteristics of the peptide sequence. Higher scores indicate a greater probability that the peptide possesses functional biological activity.
- **Instability**: This index estimates the in vitro stability of a peptide on the basis of its amino acid composition. In this study, we calculate the instability score using the **ProtParam**(Gasteiger et al. 2005), which implements the method proposed by Guruprasad et al. Peptides with an instability index below 40 are generally considered stable and suitable for experimental synthesis.

### C: Metrics for Ablation Study

To support the ablation study in this work, we introduce three custom-designed metrics that assess model performance without relying on computationally expensive structure prediction. These metrics are designed to evaluate biological relevance and diversity of the generated peptides

| Metrics(%) | ESM-2 | ProtBERT | Transformer | nn.Embedding |
|---|---|---|---|---|
| Binary Accuracy↑ | **96.03** | 93.67 | 90.20 | 75.64 |
| Top 10% Accuracy↑ | **90.18** | 87.57 | 78.20 | 39.93 |
| Top-1 Accuracy↑ | **82.51** | 78.05 | 70.23 | 20.44 |

Table 1: Performance comparison of different encoders on the protein–peptide alignment task. **Binary Accuracy** indicates whether the true peptide is ranked higher than a random baseline. **Top 10% Accuracy** measures the proportion of cases where the true peptide falls within the top 10% of the ranked candidate list. **Top-1 Accuracy** represents the percentage of samples where the correct peptide is ranked at the top (rank = 1). All models were evaluated under the same contrastive learning setup using dataset.

at the sequence level, enabling lightweight yet informative comparison across model variants. Below, we describe each metric and its computation method in detail:

**Superior Ratio** : This metric measures the proportion of test targets for which the model generates at least one peptide that matches the target protein better than the native peptide, based on dual-encoder cosine similarity. The detailed calculation procedure is described in Algorithm 1.

Algorithm 1: Calculation of Superior Ratio (SR)

**Input**: A set of test target proteins $\mathcal{T}$, trained dual-encoder model $E_{\text{prot}}$, $E_{\text{pep}}$
**Output**: Superior Ratio

Initialize counter $S = 0$
**for** each target protein $P \in \mathcal{T}$ **do**
  Generate 10 peptide sequences $\{G_1, G_2, \ldots, G_{10}\}$ for $P$ using the generative model
  Retrieve the native peptide $N$ corresponding to $P$
  Compute cosine similarity $s_{\text{native}} = \cos(E_{\text{prot}}(P), E_{\text{pep}}(N))$
  Initialize flag `has_superior` ← `False`
  **for** each generated peptide $G_i$ **do**
    Compute cosine similarity $s_i = \cos(E_{\text{prot}}(P), E_{\text{pep}}(G_i))$
    **if** $s_i > s_{\text{native}}$ **then**
      `has_superior` ← `True`
      **break**
    **end if**
  **end for**
  **if** `has_superior` is `True` **then**
    Increment $S \leftarrow S + 1$
  **end if**
**end for**
Compute Superior Ratio: $S/|\mathcal{T}|$
**return** Superior Ratio

**Inter Similarity (Inter-Sim)** : This metric quantifies the specificity of the model in generating peptides that are distinct across different target proteins. It measures the average pairwise sequence similarity between peptides generated for different targets, based on pairwise global sequence alignment scores. A lower Inter-Sim score indicates that the model produces target-specific peptides with high diversity between targets. The detailed calculation procedure is described in Algorithm 2.

Algorithm 2: Calculation of Inter Similarity (Inter-Sim)

**Input**: A set of test target proteins $\mathcal{T} = \{P_1, P_2, \ldots, P_n\}$; generative model $\mathcal{G}$
**Output**: Inter Similarity

Initialize similarity accumulator $S = 0$
Initialize pair counter $C = 0$
**for** each pair of distinct target proteins $(P_i, P_j)$ where $i < j$ **do**
  Generate 100 peptides $G_i = \{g_{i1}, g_{i2}, \ldots, g_{i100}\}$ for $P_i$ using $\mathcal{G}$
  Generate 100 peptides $G_j = \{g_{j1}, g_{j2}, \ldots, g_{j100}\}$ for $P_j$ using $\mathcal{G}$
  Initialize local similarity accumulator $s_{ij} = 0$
  **for** each peptide $g_a \in G_i$ **do**
    **for** each peptide $g_b \in G_j$ **do**
      Compute global alignment similarity $s = \text{Sim}(g_a, g_b)$
      Accumulate: $s_{ij} \leftarrow s_{ij} + s$
    **end for**
  **end for**
  Compute average similarity for this pair: $s_{ij} \leftarrow s_{ij}/(100 \times 100)$
  Accumulate global similarity: $S \leftarrow S + s_{ij}$
  Increment counter: $C \leftarrow C + 1$
**end for**
Compute final Inter Similarity: $S/C$
**return** Inter Similarity

**Intra Similarity (Intra-Sim)** : This metric quantifies the diversity of peptides generated for the same target protein by measuring the average pairwise sequence similarity within the generated peptide set. A lower Intra Similarity score indicates greater sequence diversity among peptides targeting the same protein. Specifically, for each target protein, 100 peptides are generated by the model, and the global alignment similarity is calculated for every unique peptide pair. The average similarity across all pairs is computed per target, and the final Intra Similarity score is obtained by averaging over all targets. The detailed calculation procedure is provided in Algorithm 3.

| Methods | Intra-Sim↓ | Inter-Sim↓ | Superior ratio↑ | GACD↓ | Bioactivity↑ | Instability↓ |
|---|---|---|---|---|---|---|
| PepCCD (w/o Align & Pre-training) | 0.3532 | 0.4097 | 21.05 | 0.3729 | 0.2720 | 57.68 |
| PepCCD (w/o Pre-training) | 0.3347 | 0.3670 | 37.79 | 0.2804 | 0.3439 | 52.94 |
| PepCCD (w/o Align) | 0.2845 | 0.3513 | 22.96 | 0.2418 | 0.3383 | 44.63 |
| PepCCD | 0.2392 | 0.2445 | 44.97 | 0.1754 | 0.4419 | 37.66 |

Table 2: Ablation study results of PepCCD and its three variants across six evaluation metrics. Note that Intra Similarity measures the sequence similarity among peptides generated for the same target, reflecting diversity; Inter Similarity measures the similarity between peptides generated for different targets, reflecting the model's ability to distinguish between targets; Superior ratio evaluates the target-specificity of the generated peptides.

| Methods | Type | ipTM(best)↑ | ipTM(avg)↑ | RT-score(best)↓ | RT-score(avg)↓ |
|---|---|---|---|---|---|
| RFdiffusion | Structure | **0.7838** | **0.5960** | **-394.422** | **-242.238** |
| PepPrCLIP | Sequence | 0.7141 | 0.4901 | -188.933 | -157.573 |
| PepCCD | Sequence | <u>0.7418</u> | <u>0.5557</u> | <u>-363.481</u> | <u>-188.933</u> |

Table 3: Performance on target-oriented peptide design. In the column headers, **(best)** denotes the best single peptide designed for each target, whereas **(avg)** denotes the average performance across all targets.

Algorithm 3: Calculation of Intra Similarity (Intra-Sim)

**Input**: A set of test target proteins $\mathcal{T} = \{P_1, P_2, \ldots, P_n\}$; generative model $\mathcal{G}$
**Output**: Intra Similarity

1: Initialize similarity accumulator $S = 0$
2: **for** each target protein $P_i \in \mathcal{T}$ **do**
3: Generate 100 peptides $G_i = \{g_{i1}, g_{i2}, \ldots, g_{i100}\}$ for $P_i$ using $\mathcal{G}$
4: Initialize local similarity accumulator $s_i = 0$
5: Initialize pair counter $c_i = 0$
6: **for** each pair of distinct peptides $(g_a, g_b)$ in $G_i$ where $a < b$ **do**
7: Compute global alignment similarity $s = \text{Sim}(g_a, g_b)$
8: Accumulate: $s_i \leftarrow s_i + s$
9: Increment pair counter: $c_i \leftarrow c_i + 1$
10: **end for**
11: Compute average similarity for $P_i$: $s_i \leftarrow s_i / c_i$
12: Accumulate global similarity: $S \leftarrow S + s_i$
13: **end for**
14: Compute final Intra Similarity: $S / |\mathcal{T}|$
15: **return** Intra Similarity

The three evaluation metrics are well-aligned with the core objectives of our task, which involve generating peptides that are both target-specific and diverse. Specifically, Intra Similarity captures the diversity of generated peptides for the same target; Inter Similarity reflects the model's ability to distinguish between different targets; and Superior Ratio directly evaluates the target-specificity of generated peptides compared to native binders. As shown in Table 2, the ablation study demonstrates that each individual component of PepCCD plays a vital role in enhancing both the specificity and diversity of the generated peptides, validating the effectiveness of the overall architecture design.

### D: Details of GACD Calculation

We specifically designed a metric to quantify the overall compositional difference between generated peptides and native template peptides in terms of amino acid use, called the global amino acid composition discrepancy **(GACD)**. This metric is calculated based on global frequency distributions of the 20 standard amino acids and measures the Euclidean distance between the distributions of the generated peptide set and the reference template set. A lower GACD score indicates that the amino acid composition of the generated sequences matches more closely that of the natural peptides. The computation is defined as follows:

$$\text{GACD} = \sqrt{\sum_{i=1}^{20} \left(f_{\text{template}}\left(a_i\right) - f_{\text{generated}}\left(a_i\right)\right)^2} \quad (1)$$

where $f(a_i)$ represents the frequency of the amino acid. Specifically, we collect all peptides generated by the model and all native template peptides separately, compute the global amino acid frequency distribution for each set, and then calculate the Euclidean distance between these two distributions using the formula above to quantify the overall compositional discrepancy between generated and native sequences.

### E: Detailed Comparison of ipTM Scores Across Peptide Design Models

In the main text, we compared the interaction capabilities between the generated peptides and target proteins across three models, as shown in Table 3. To extend this analysis, we further utilized AlphaFold 3 to predict complex structures and compute the ipTM(avg) score. Using the ipTM score of the corresponding native template peptide as a reference, we defined the hit rate as the proportion of generated peptides with ipTM(avg) $\geq$ template ipTM, serving as a metric to evaluate the model's ability to generate high-quality structures.

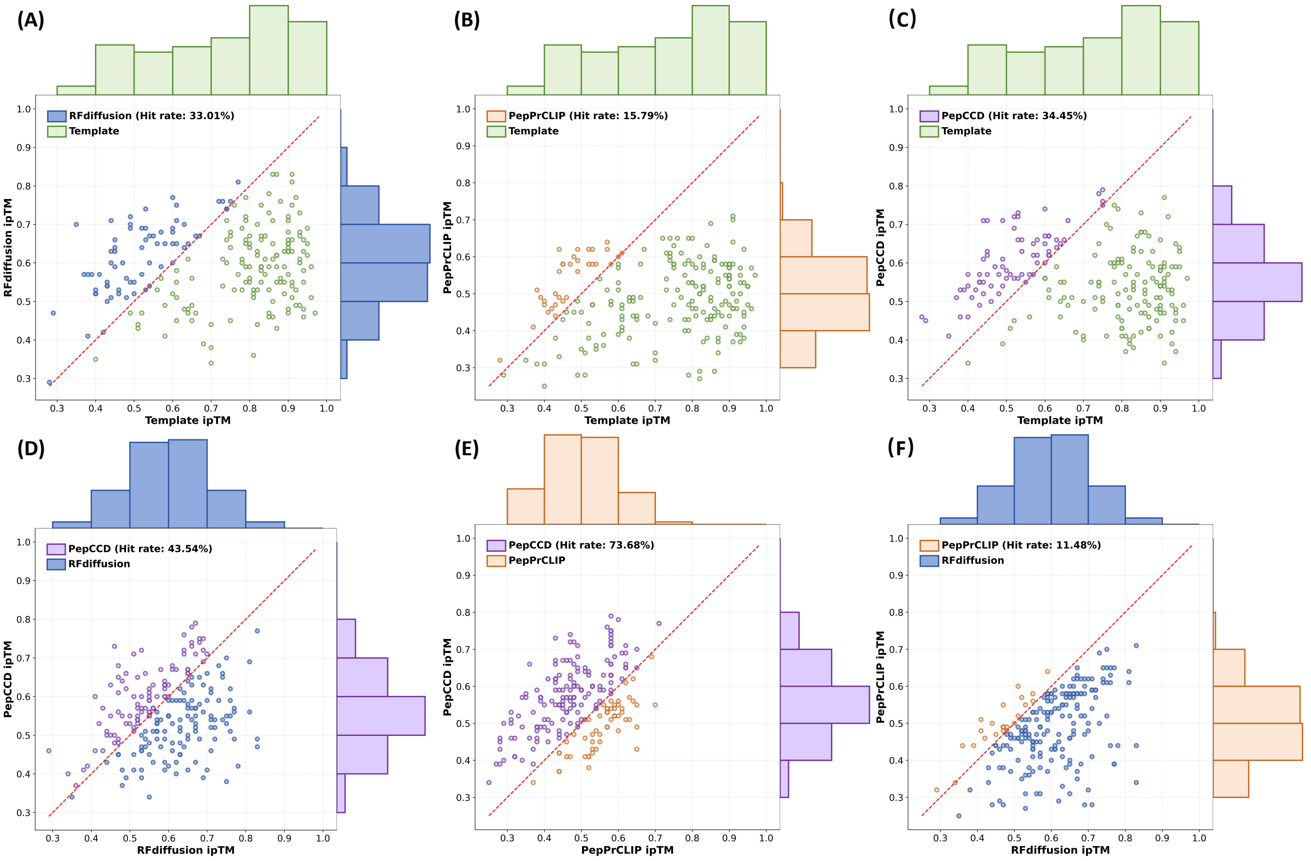


Figure 1: Comparison of ipTM(avg) scores across different models and model-to-model hit rate evaluations. (A)–(C): Distribution of ipTM(avg) scores of peptides generated by RFdiffusion, PepCCD, and PepprCLIP, respectively, each compared against their corresponding native template peptides. (D)–(F): Cross-model hit rate analysis, where each subplot shows the proportion of peptides from one model achieving equal or higher ipTM(avg) scores than those from another model, used as the reference template.

Interestingly, although PepCCD exhibits a slightly lower average ipTM(avg) score compared to RFdiffusion, it achieves a marginally higher hit rate. To gain deeper insights into the differences among these models, we further used the ipTM(avg) values generated by each of the three models as "template references" and evaluated the hit rate performance of the other two models accordingly. As shown in Figure 1, when using RFdiffusion—whose ipTM scores are the highest—as the reference template, PepCCD achieves a hit rate of 43.54%, significantly outperforming PepprCLIP, which achieves only 11.48%. From Figure 1(D), a moderate positive correlation can be observed between PepCCD and RFdiffusion across multiple targets. This suggests that, despite their fundamentally different modeling strategies, PepCCD is still able to partially capture important conformational patterns that are typically driven by structural context.

Furthermore, when using the sequence-based PePprCLIP as the reference template, the hit rate of PepCCD significantly increases to 73.68%, as shown in Figure 1(E). This clearly demonstrates that, under the same sequence-guided conditions, PepCCD exhibits stronger generative stability and target recognition capability. In summary, PepCCD does not rely on a few high-scoring samples to boost its average performance; rather, it generates a more concentrated distribution of peptides that can consistently interact with the target protein. This also explains why PepCCD achieves the best performance when using native template peptides as references.

### F: Details of Molecular Dynamics Simulations

**System Construction and Preprocessing** We selected the receptor-binding domain (RBD) of the SARS-CoV-2 spike protein (PDB ID: 6M0J(Lan et al. 2020)) as the receptor for constructing peptide–protein complexes using the candidate peptides generated by each model. For each method, 10 representative peptides were selected. The peptide structures were predicted using AlphaFold3, and docking was performed using Rosetta FlexPepDock to ensure that the peptides were positioned near the functional binding pocket between the RBD and ACE2(Yan et al. 2020). Docking conformations for each complex were evaluated using

| Methods | Type | MM/GBSA↓ | MM/PBSA↓ |
|---|---|---|---|
| RFdiffusion | Structure | -18.24 ± 16.72 | -18.91 ± 12.06 |
| PepPrCLIP | Sequence | -25.09 ± 14.97 | **-27.02 ± 10.31** |
| PepCCD | Sequence | **-25.11 ±6.88** | -26.31 ± 6.56 |

Table 4: Binding free energy comparison of peptides designed by different methods. Average binding free energies (in kcal/mol) were calculated using MM/GBSA and MM/PBSA methods based on 100 ns molecular dynamics simulations. Lower values indicate stronger binding affinity. Bold values represent the best performance, while underlined values indicate the second-best.

the score_jd2 module. The top-scoring conformation with a physically reasonable structure was selected for downstream molecular dynamics simulations.

**Force Field and Simulation Parameters** All molecular dynamics simulations were performed using the GROMACS 2024.3 software package(Abraham et al. 2015). Initial peptide–protein complex systems were constructed via the CHARMM-GUI platform(Jo et al. 2008). Given the presence of glycosylation in the receptor-binding domain (RBD) of the SARS-CoV-2 spike protein, we employed the **Glycan Reader & Modeler** module of CHARMM-GUI to ensure accurate modeling of glycosylation. The CHARMM36m force field was used for all parameterizations. TIP3P was chosen as the water model, and $Na^+$ and $Cl^-$ ions were added to reach a physiological salt concentration of 0.15 mol/L.

The simulation workflow consisted of three major stages: energy minimization, equilibration, and production runs. Parameters for each stage were based on default configuration files provided by CHARMM-GUI to ensure consistency and stability of the simulated systems under near-physiological conditions.

**Binding Free Energy Calculation** To evaluate the binding affinity between the designed peptides and the target protein, we performed binding free energy calculations based on molecular dynamics trajectories. The **gmx_MMPBSA** tool was employed to estimate the binding free energies using both the MM/GBSA and MM/PBSA methods(Genheden and Ryde 2015). A total of 1000 frames were extracted from the 100 ns production trajectories at 10 ps intervals. The energy calculations were carried out using the same force field (CHARMM36m) and solvent model (TIP3P) as in the simulations. The system was maintained under a physiological salt concentration of 0.15 mol/L throughout the calculations. Final outputs included total binding free energies and their respective components, facilitating comparative analysis of peptide binding stability across different design models.

As shown in Table 4 and Figure 2, the sequence-guided models exhibit the best overall performance in molecular dynamics simulations. Among them, PepCCD consistently achieves the lowest average binding free energies across MM/GBSA calculation. Moreover, it demonstrates the smallest standard deviation in energy values, indicating superior stability and consistency of peptide–protein interactions throughout the simulation trajectory. These results suggest that PepCCD not only generates peptides with strong binding affinity but also maintains reliable structural integrity under dynamic physiological conditions. Notably, although RFDiffusion performs well on static target-binding metrics, it demonstrates the poorest performance in molecular dynamics simulations. This discrepancy may stem from the fact that the selected complex structure (PDB: 6M0J) lacking a native peptide ligand for the RBD, which prevents RFDiffusion from fully leveraging the advantages of its pre-trained structural information.

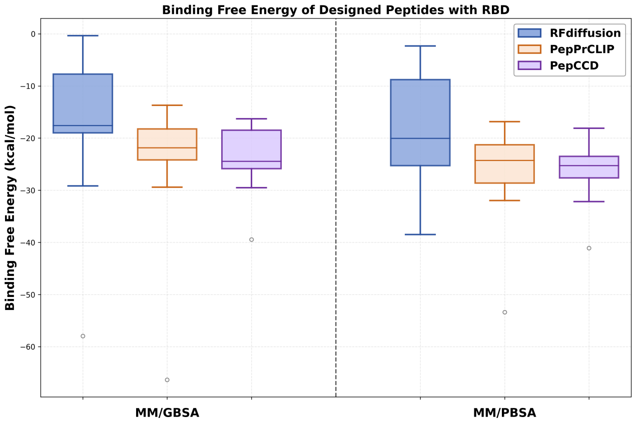


Figure 2: Based on the simulation trajectories, the binding free energies of each peptide–protein complex were calculated using the MM/GBSA and MM/PBSA methods, in order to assess the binding affinity between the peptides and the target protein.

### G: Hyperparameter Settings for PepCCD

The training process of PepCCD is divided into three sequential stages. In the first stage, a small learning rate of 3e-5 and a batch size of 16 are used for 50 epochs to warm up the model and stabilize parameter updates. In the second stage, the learning rate is increased to 5e-4, the batch size remains at 16, and a diffusion process with 500 timesteps is introduced. This stage is trained for 50,000 steps, allowing the model to fully learn the peptide generation patterns. The third stage maintains the same learning rate and diffusion timesteps but reduces the number of training epochs to 10,000, aiming for fine-tuning and convergence. The batch size is kept constant across all stages.
